# Flux Sampling in Genome-scale Metabolic Modeling of Microbial Communities

**DOI:** 10.1101/2023.04.18.537368

**Authors:** Patrick E. Gelbach, Stacey D. Finley

**Affiliations:** Alfred E. Mann Department of Biomedical Engineering, University of Southern California, Los Angeles, CA 90089, USA; Department of Quantitative and Computational Biology, University of Southern California, Los Angeles, CA 90089, USA; Mork Family Department of Chemical Engineering and Materials Science, University of Southern California, Los Angeles, CA 90089, USA

## Abstract

Microbial communities play a crucial role in ecosystem function through metabolic interactions. Genome-scale modeling is a promising method to understand these interactions. Flux balance analysis (FBA) is most often used to predict the flux through all reactions in a genome-scale model. However, the fluxes predicted by FBA depend on a user-defined cellular objective. Flux sampling is an alternative to FBA, as it provides the range of fluxes possible within a microbial community. Furthermore, flux sampling may capture additional heterogeneity across cells, especially when cells exhibit sub-maximal growth rates. In this study, we simulate the metabolism of microbial communities and compare the metabolic characteristics found with FBA and flux sampling. We find significant differences in the predicted metabolism with sampling, including increased cooperative interactions and pathway-specific changes in predicted flux. Our results suggest the importance of sampling-based and objective function-independent approaches to evaluate metabolic interactions and emphasize their utility in quantitatively studying interactions between cells and organisms.

## 2. Introduction

Microbes are essential components of all living ecosystems, and the metabolic interactions between them are a significant factor in the functioning of these ecosystems. Microbe-microbe metabolic interactions affect nutrient cycling, energy production, and the maintenance of microbial diversity^1–3^. Though our understanding of those microbial communities is aided by metagenomics and *in vitro* analyses, there is a significant gap in mechanistic understanding of the makeup and interactions between members of microbial consortia^4,5^.

Genome-scale modeling has emerged as a promising method by which we can probe an organism’s metabolic states, behaviors, and capabilities, alone or as a community^6–12^. Genome-scale metabolic modeling is a mathematical approach that uses the known biochemical reactions of a species to reconstruct a genome-scale metabolic network. Genome-scale models (GEMs) provide a holistic view of an organism’s metabolism, allowing for mathematical analyses that simulate metabolic fluxes and thus provide insight into metabolic pathways and physiological processes. The genome-scale model consists primarily of a stoichiometric matrix, characterizing the interconversion of metabolites by the set of metabolic reactions, linked with a set of Boolean expressions describing the gene-protein-reaction relationships^40^. Flux balance analysis is a constraint-based approach for analyzing that metabolic network to predict metabolic fluxes through the GEM.

Much work has recently been applied to understand the metabolic interactions of a microbial community in various contexts, including the human gut microbiota and in environmental bioremediation^14–19^. Given the ubiquity of microbial activity, there is substantial value in using metabolic modeling to understand these communities’ emergent behaviors and abilities.

Most metabolic modeling of microbial interactions is performed in one of three ways (**Figure 1**): (1) *compartmentalization*, wherein two metabolic models are merged into a single stoichiometric matrix with a shared compartment representing the extracellular space, (2) *lumped model* (also called “enzyme soup”) approach, where all metabolites and reactions are pooled into a single model in proportion to the community makeup, and (3) *costless secretion*, where models are separately simulated while dynamically and iteratively updating the simulated environment by adjusting the models’ exchange reactions and available nutrients based on metabolites that can be secreted without decreasing growth (costless metabolites)^19–26^.

**Figure 1:**
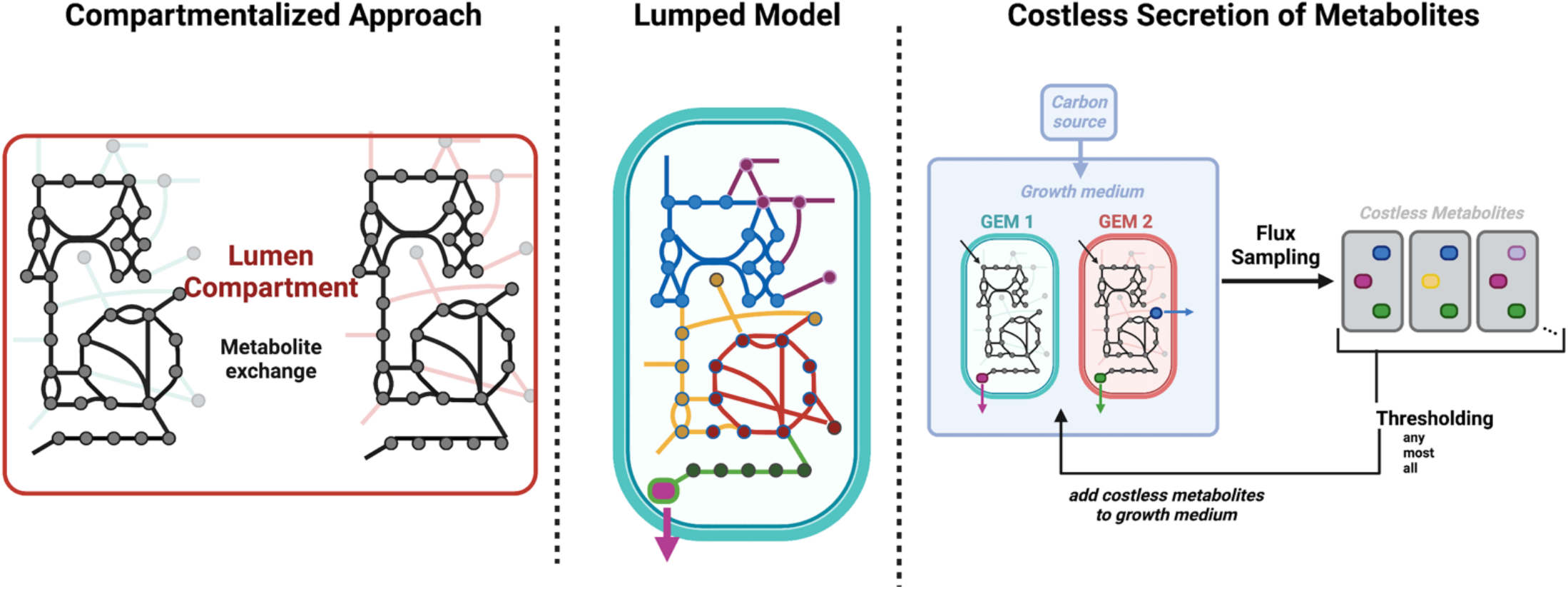
Approaches for genome-scale metabolic modeling of communities. Metabolic modeling of microbial communities is largely performed using (A) Compartmentalization: a single stoichiometric matrix representing the two models joined by a lumen compartment wherein metabolites can be freely exchanged; (B) Lumped model: a single stoichiometric matrix representing the union of each model’s reactions, thereby ignoring all separation between cells; and (C) Costless secretion: individual stoichiometric networks for each model, whose exchange reactions are constrained to reflect the shared extracellular media.**1**

Each of these approaches has shown promise, and selection of which approach to use heavily depends on available data and models and the intended goal of the analysis. As currently implemented, each method uses flux balance analysis (FBA), a linear programming technique that predicts the flow of material through the metabolic network^27–29^. FBA depends on the maximization of an objective function, and maximizing biomass production is most commonly used. Optimizing for biomass assumes species are entirely oriented towards maximal growth, thus ignoring the multiplicity of achievable sub-optimal phenotypes^30^. When simulating the metabolism of a community, this assumption can disregard the variety of metabolic interactions that the microbes may carry out. Furthermore, the selection and definition of the best objective function substantially affect model predictive power and generated results^31–34^.

As an alternative to FBA, flux sampling has recently been used to predict flux distributions in a variety of cases and may provide a more holistic and accurate description of the cell’s flux distribution^35–39^. This is done by randomly generating many flux values for each reaction in a genome-scale metabolic model, while respecting its defined constraints, such as mass or energy balance and thermodynamic restrictions. Flux sampling employs Markov chain Monte Carlo methods to estimate cellular flux and generate many feasible metabolic flux distributions. Flux sampling estimates the most probable network flux values, enabling statistical comparisons of the flux distributions. Notably, the approach does not require a selected cellular objective, thus reducing user-introduced bias on model predictions and exploring the entire constrained solution space. The approach therefore enables studies of phenotypic heterogeneity, as a single constrained model can generate a range of flux predictions. However, flux sampling has not been widely employed in analyses of microbial communities. Furthermore, comparisons between FBA-based and sampling-based analyses of communities are currently lacking.

In this work, we apply flux sampling to existing analyses of microbial metabolic interactions, showing the range of potential consortia-wide flux distributions achievable with genome-scale modeling. We find significant differences in model predictions between FBA and flux sampling, with substantial heterogeneity across sampled simulations. We see emergent patterns at sub-maximal growth rates, such as increased cooperation between microbes in anoxic conditions compared to oxygen-rich environments. In total, we systematically evaluate the effect of flux sampling, and emphasize the utility of objective function-agnostic approaches to evaluate metabolic interactions.

## 3. Methods

### 3.1 GEMs

Magnusdottir *et al*. generated the AGORA dataset, a collection of 773 (and later, 7206 in AGORA2) genome-scale metabolic models comprising the human gut microbiome. These models were simulated to understand their metabolic behavior when grown in pairwise combinations, using the approach developed by Kiltgord and Segre^16,42,43^. Notably, the analysis constrained the models with distinct *in silico* diets and aerobic states.

We randomly selected 75 of the AGORA models, analyzed all unique pairwise combinations (2775), and implemented three distinct approaches to study metabolic interactions between microbes. In this way, we demonstrate the utility of each approach, compared to flux sampling, while limiting computational intensity.

### 3.2 Flux sampling

We use the Constrained Riemannian Hamiltonian Monte Carlo (RHMC), which has recently been shown to be substantially more efficient than prior sampling algorithms^41^.

### 3.3 Compartmentalization

The pairwise interaction approach used by Magnusdottier and coworkers is as follows^17^:

**Step 1:** Select two models.

**Step 2:** Introduce the lumen compartment, which joins the two models into a merged model, where the two microbes can secrete and uptake metabolites.

**Step 3:** Constrain the model by adjusting exchange reaction bounds to reflect the chosen diet and extracellular conditions.

**Step 4:** Simulate monoculture by shutting off one of the two models by inactivating all its reactions (setting the reaction upper and lower bound to 0 flux). Then simulate the active individual model by optimizing for growth.

**Step 5:** “Shut off” the second individual model by inactivating all its reactions. Then simulate the active individual model by optimizing for growth.

**Step 6:** Restore the activity of both individuals in the merged model and optimize each microbes’ objectives separately. This predicts growth while allowing the exchange of metabolites across the lumen, simulating co-culture.

**Step 7:** Compare paired growth with the individual growth simulations of steps 4 and 5. If paired growth was 10% higher or lower than individual growth, the model was considered to grow faster or slower, respectively, in co-culture than alone.

We replaced the FBA optimization in steps 4, 5, and 6 with flux sampling as an alternative way to predict cellular flux. We used the RHMC algorithm and generated 1000 flux distribution samples at each step. We, therefore, had a range of reaction fluxes (including growth rates) for both microbes, in mono- and co-culture, with and without oxygen, and with two different simulated diets (Western and High Fiber). We then categorized all possible combinations of sampled growth rates, following previously described classes: parasitism, commensalism, neutralism, amensalism, competition, or mutualism^17^. For example, if both models grew more in co-culture than alone, the interaction was classified as mutualism.

We also identified distinct interaction regimes between the two microbes by ordering the sampled growth rates. That is, we found the range of different interaction types as a function of the different growth rates (and thus, growth demands). We note that the interaction regimes predicted here differ from the Pareto analysis performed by Magnusdottir et al., as calculation of the Pareto front relies on biomass optimization with FBA while iteratively updating and fixing growth rates for each model^16^.

### 3.3 Lumped model

Blasco *et al*. extended the AGORA set of metabolic models by adding degradation pathways that allow for the simulation of the effect of many human diets on the activity of the gut flora^26^. After adding those metabolic reactions involved in degradation, they merged all individual microbe models into a supra-organism model. By pooling all GEMs, they made a single lumped model comprising all metabolic reactions and metabolites in the population.

This process is often called a “mixed-bag” or “bag-of-genes” approach and is the simplest form of genome-scale modeling of bacterial communities. It does not assume any spatial or temporal separation between the species and involves the consolidation of ubiquitous metabolic reactions^44–47^. Nevertheless, the approach has been effective at predicting the metabolic behavior of consortia while minimizing computation time and reducing model size.

The authors used flux variability analysis (FVA) to identify and correct blocked or low-confidence reactions and identify the microbial metabolic byproducts produced by the microbiota’s fermentation of lentils. However, the model was not simulated to predict species growth within the community. We, therefore, applied the model to predict consortia behavior. With “mixed-bag”, lumped modeling of metabolism, it is common to either merge all individual model biomass reactions into a supra-organism growth equation or, as chosen by Blasco et al., to maintain each model’s biomass reaction within the pooled network. That allows for the prediction of each microbe’s growth.

We generated flux samples of the lumped model and compared them to the case where each species’ biomass reaction is optimized alone and to the case where the population’s overall growth is maximized. We calculated the “optimal community growth rate” by finding the maximum growth rate possible for all models simultaneously and setting all biomass reactions’ lower bound values to that flux value.

### 3.4 Costless Secretion

Pacheco et al. showed that the secretion of costless metabolites (species that are freely secreted as the byproduct of a cell’s metabolism, without inhibiting fitness) are critical drivers of the metabolic interactions between cells^48^. The approach is a quasi-dynamic method, as it maintains the modeling assumption that the system is at a steady state but successively updates the environment shared by the two simulated cells. The method calculates the growth of each model at each simulation iteration, finds the secreted metabolites, then updates the simulated media until the media is stable.

The costless growth approach is as follows:

**Setup:** Select a simulated minimal media definition (i.e., DMEM, M9, SSM, etc.), and define the metabolites that comprise that medium. Select *N* models to be simulated (i.e., two models for pairwise interactions, three models to simulate a community of three microbes, etc.) Select *M* metabolites to be individually or provided in addition to the minimal media (list the carbon sources to be provided, and choose m to be given at a time). Define whether the model will be simulated in an aerobic or anaerobic environment.

**Step 1:** Simulate a minimal media condition by setting all exchange reactions’ upper bound values to 0 unless the reaction exchanges metabolites contained in the media.

**Step 2:** Provide carbon source(s) *M* by setting the upper bound(s) of exchange reaction(s) for the corresponding metabolite(s) to be unconstrained.

**Step 3:** Simulate the models by optimizing for growth.

**Step 4:** Note the resulting flux values for the models’ transport reactions; if a metabolite is predicted to be exported into the media, then explicitly add that metabolite to the simulated media (again, by adjusting the models’ upper bound for that metabolite’s import reaction).

**Step 5:** Repeat steps 3-4 until no additional metabolites are secreted, arriving at the simulation’s final predicted growth rates.

Steps 1-5 can be repeated for a different carbon source (or combination of carbon sources) to be added to the simulated media.

We adjusted step 3, replacing FBA optimization with flux sampling using the RHMC algorithm, simulating pairwise growth with a single metabolite source. Because the costless secretion approach repeats the model simulation steps until media convergence, we introduced a thresholding term to consolidate the results from each simulation round. In particular, we define the set of secreted metabolites (and thus update the extracellular media) based on whether all, most, or any of the sampled flux distributions show a metabolite is secreted. For example, if metabolite *M* were secreted in 200 of the 1000 generated flux distributions, it would only be added to the extracellular media in the “any” cutoff simulation for the next round. If secreted in 750 of the 1000 distributions, it would be added to the “any” and “most” analyses. For the “all” cutoff, that metabolite would only be added to the media for the following iteration if all 1000 flux distributions showed that metabolite was secreted.

Thresholding is currently required because of the computational time needed for flux sampling. Without thresholding at each simulation, the number of sampled points needed will increase exponentially with each round of expansions. We use each cutoff to demonstrate and assess the introduced variability, allowing us to compare the outputs with each set threshold.

## Results

### 4.1 Compartmentalized modeling

Magnusdottir et al. developed AGORA (later updated as AGORA2), a resource for the semi-automated generation of genome-scale metabolic models. They applied the set of models to predict the pairwise interactions between microbes, showing how the individuals’ metabolic potential drives the emergent behavior of the pair. However, their predictions of metabolic activity assume that each microbe is oriented toward achieving maximal growth. We introduced flux sampling to the pairwise simulation framework, thus permitting any flux distribution from the confined flux space to be included in the assessment of metabolic interactions. We selected 50 individual models and paired each together, generating 2775 unique pairwise analyses that spanned the range of microbial taxa in the AGORA dataset. Each paired model was sampled 1000 times, with and without metabolite exchange between the microbes being allowed. As described in the Methods, the interaction type was calculated for the anaerobic and aerobic states with two unique simulated diets.

By analyzing the most common interaction for each pair, we calculated the total percentage of each interaction type, shown in Figure 2A. As expected, slight differences exist between the optimization and sampling-based analyses. Namely, antagonistic interactions (competition, amensalism, and parasitism) tend to make up a smaller percentage of the entire set with sampling instead of optimization (61% compared to 74%). There is an increase of 11% in positive and net-neutral interactions (commensalism, neutralism, and mutualism) with sampling compared to optimization-based analysis. Cases of neutralism increased from 6% to 18%, and frequency of mutualism increased from 7% to 13%. The increase in cooperation is particularly prevalent with anaerobic analyses, from 30% to 44%. Previous work has highlighted that anoxic conditions induce mutualism; this effect is notably amplified with sampling compared to simulations maximizing biomass^23^. When sampling the possible fluxes of anaerobic conditions, there is a higher frequency of non-inhibitory relationships. In particular, there is a substantial reduction in parasitism with a nearly equivalent increase in neutralism.

**Figure 2:**
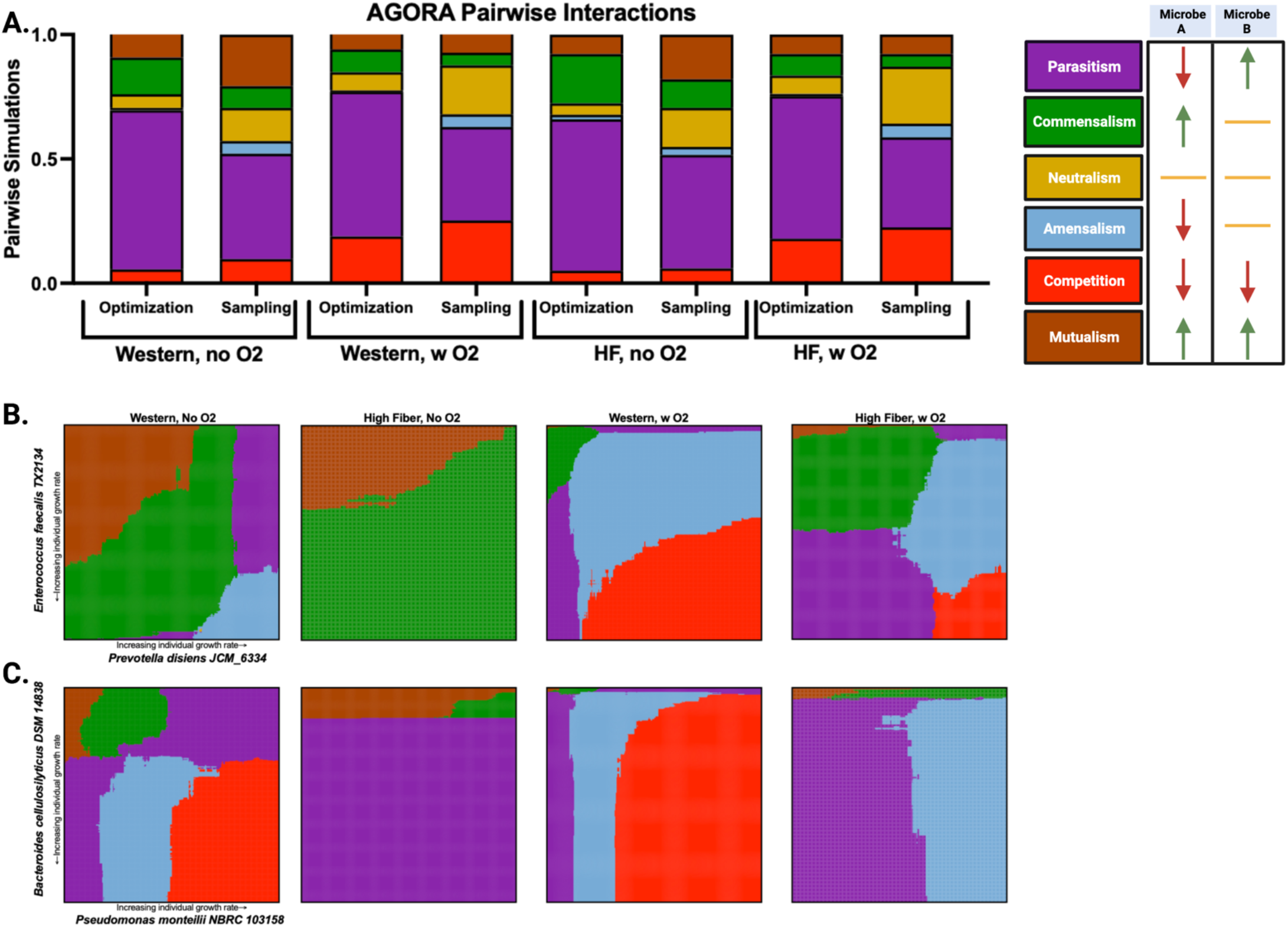
Pairwise analyses of the AGORA/AGORA2 set of models.(A.) We simulated 2775 pairs of metabolic models on two simulated diets with and without the presence of oxygen and calculated the expected interaction type. Interactions are defined and colored according to the labels on the far right. Expected interaction type when pairing enterococcus faecalis and prevotella disiens (B) and bacteriodes celiilosilyticus and pseudomonas montelli (C) and sampling a range of growth rates.

Furthermore, we see an increase in symmetrical interactions (mutualism, neutralism, and competition, from 25% to 47%), suggesting that without orienting all metabolism towards optimal growth, a community of bacterial species may be more inclined towards population stability. This is because abundances tend to remain steady when primarily exhibiting those three interaction motifs^49,50^. Notably, the trends of increased anoxic cooperation and symmetrical interactions are found irrespective of diet constraints. This suggests that the submaximal predicted growth rate allowed with sampling, and not the models themselves or extrinsic factors (such as the nutrients provided), is driving the observed outcomes.

Because flux sampling gives a distribution of growth rates and the corresponding flux distributions providing for that growth, it is possible to calculate the expected pairwise interactions likely for each combination of individual growth rates. We ranked the sampled growth rates for each microbe and calculated the most commonly predicted paired growth rate, thus giving interaction types for each growth rate. The sampling-based approach highlights the variety of interactions possible between two microbes, especially given variation in simulated conditions. Figure 2 shows this analysis for two sets of paired example microbes, similar to chemical phase diagrams. The x- and y-axes represent the individual sampled growth rates, and their intersection is colored according to the most likely expected metabolic interaction motif. Figure 2B shows the pairwise interactions of *enterococcus faecalis TX2134*, a gram-positive nonmotile microbe, and *prevotella disiens JCM 6632*, a gram negative bacilii-shaped bacterium. Figure 2B shows the interactions of *bacteriodes celiilosilyticus* and *pseudomonas montelli*, two gram-negative and rod-shaped microbes. We show these calculations for four distinct extracellular conditions: Western and high fiber diets, with and without oxygen.

When simulating the interactions of the *Enterococcus* and *Prevotella* strains in Figure 2B, five distinct types of interactions are possible, depending on the simulated environment and each species’ growth rate. Anaerobic states (columns 1 and 2) show a predominance of commensalism or mutualism, though parasitism and amensalism are expected when *Prevotella* is rapidly proliferating. In the presence of oxygen (columns 3 and 4), there are several diet-independent trends: low growth of both microbes causes commensalism; high *Enterococcus* and low *Prevotella* growth rates cause parasitism; low *Enterococcus* and high *Prevotella* growth rates cause amensalism; and high growth of both causes competition. At intermediate growth rates, the effect of diet is more apparent, as Western diet constraints drive amensalism and high fiber constraints push the interaction toward commensalism, parasitism, or amensalism.

Similar insights can be gained when analyzing the interactions between *Bacteroides* and *Pseudomonas*. For example, the anoxic high fiber condition is relatively invariant, as the two microbes show parasitism at nearly all individual growth rates. Alternatively, there is a large set of potential outcomes when simulating an anoxic state with a Western diet; the individual microbe growth rates can elicit widely distinct interaction motifs. It is possible to see a single microbe “dominate” or drive the observed interaction: in the oxygen-rich simulations, changes in *Pseudomononas* growth determine the outcome, largely irrespective of a changing *Bacteriodes* growth rate.

Similar analyses can be performed for all combinations of models. In sum, this sampling-based approach highlights the variety of interactions possible between two simulated microbes, especially given variety in modeled conditions.

### 4.2 Lumped model

By pooling metabolic reactions, it is possible to generate a single GEM that represents community metabolic activity. The lumped GEM can then be analyzed using the same constraint-based approaches typically utilized for single-species models. Though the technique removes all separation between microbes, it can be a useful approach for assessing the activity and potential of the community. However, all analyses of such “bag of genes” or “enzyme soup” approaches have explicitly assumed that the community aims to maximize growth by assigning an objective function. No studies have assessed the effect of flux sampling on the community metabolic state. We selected the AGREDA pooled model, which combined 538 AGORA models into a single metabolic network. We analyzed the lumped model with three distinct approaches: (1) iteratively setting each individual’s biomass reaction as the objective and then solving the FBA problem (optimization), (2) performing flux sampling on the network’s flux solution space (flux sampling), and (3) finding the maximal rate at which all microbes can simultaneously grow, then sampling the solution space when that value is set as the lower bound for each microbe’s growth rate (termed an “optimal community”). These analyses allowed us to compare the flux distributions achieved through FBA, flux sampling, and flux sampling of the “best state” of the microbial community.

When comparing flux sampling of the network with FBA, we first assessed the variation between pathway fluxes to identify large-scale metabolic shifts. Interestingly, flux sampling was not equally influential across all pathways but disproportionately affected particular subsystems. Figure 3A shows the median normalized flux value through each pathway predicted by sampling (y-axis) and the median flux value through the pathway when individually optimizing each of the 531 biomass reactions in the model (x-axis). That suggests the parts of the network that may be more influential and impactful in community activity when separated from the requirement of maximizing cellular growth. Notably, thiamine metabolism, terpenoid backbone synthesis, tannin degradation, pyrimidine synthesis, and NAD metabolism saw substantially higher fluxes in the sampling approach. A similar plot comparing the community-constrained sampling with the FBA approach is in the supplement, in Figure S1. For example, we plot the individual fluxes through the NAD metabolism subsystem obtained by the three techniques used to predict reaction flux (Figure 3B). Maximization of biomass causes consistently low pathway flux, while unconstrained or optimal community-constrained sampling predicts a broader range of flux values that are higher compared to the fluxes predicted when biomass production is maximized. The case with optimal community-constrained sampling produces slightly elevated pathway flux compared to unconstrained sampling.

**Figure 3:**
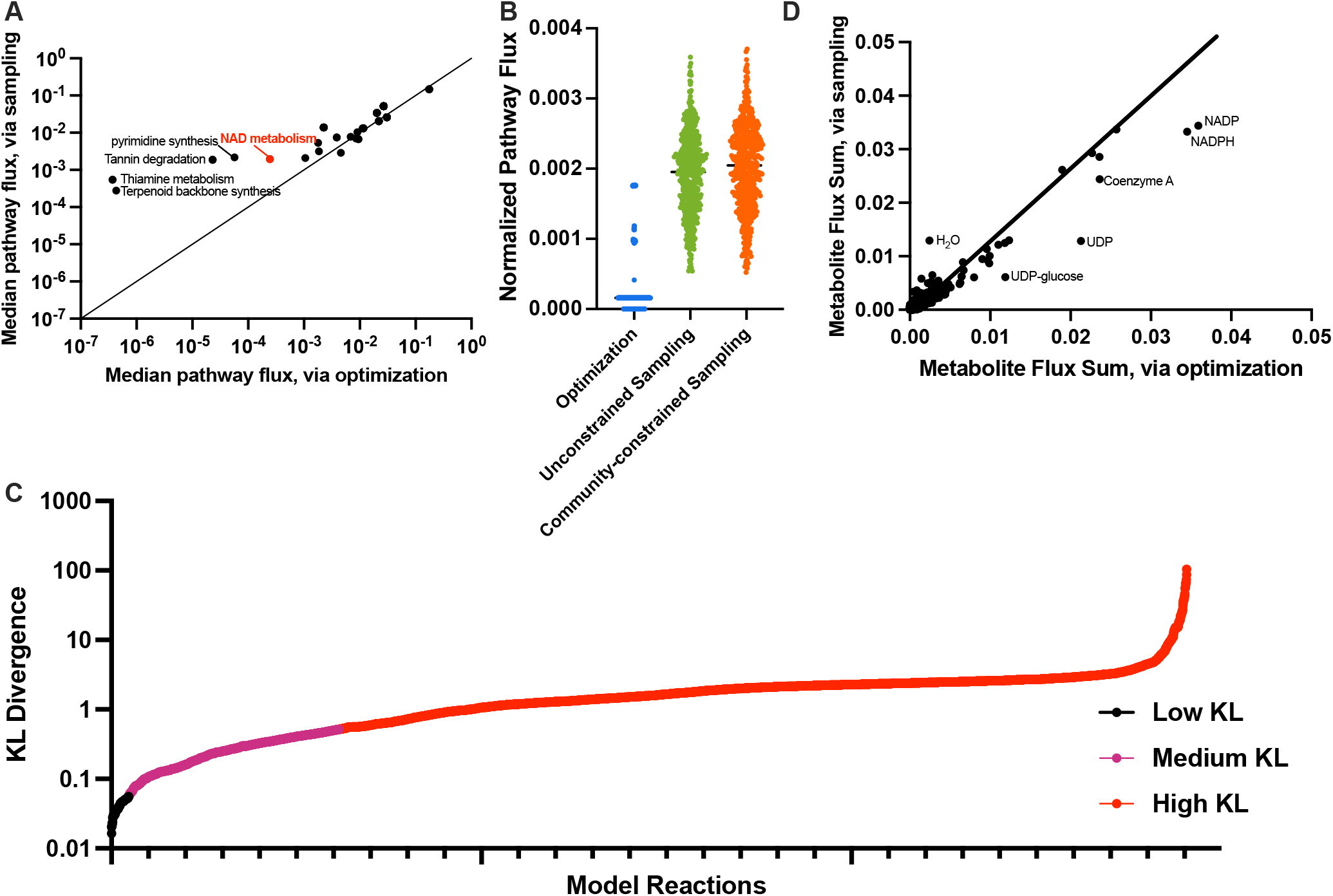
Pooled Model Analyses. (A) Median pathway flux values predicted by unconstrained flux sampling compared to optimization of biomass. Subsystems that have significantly different median fluxes are labeled. (B) Reaction fluxes for the NAD metabolism pathway predicted with each technique. (C) KL divergence between the distribution of fluxes achieved via optimization and sampling. (D) Comparison of the flux-sum value for each metabolite for unconstrained flux sampling and optimization of biomass.

At the reaction level, differences emerge between the three community analysis methods. We compared the flux distributions for each reaction, calculating the bi-directional KL divergence values^51^, shown in Figure 3C. We classified the difference in the median fluxes for each reaction for two analysis methods into low, medium, and high divergence categories. This calculation revealed that very few (76 of the 5499 reactions, to the left of the leftmost vertical line) show close alignment between unconstrained flux sampling and optimization of biomass^37,39^. The predicted flux through most reactions (86%, shown to the right of the second vertical line) is “widely divergent” between the two approaches. This points to the substantial differences between the optimal growth state and the total solution space. Interestingly, a metabolite-centric view based on metabolite flux-sum analysis shows similar turnover rates for the metabolites across the two analysis methods (Figure 3D). Metabolites that vary widely between the two conditions include NADP, NADPH, coenzyme A, UDP, and UDP-glucose (higher with optimization) and water (higher with sampling).

### 4.3 Costless secretion

Pacheco et al. argued that the secretion of “costless” metabolites (byproducts of the cell’s metabolism that are released without causing a loss of fitness) might be a primary driver of interspecies interactions within a microbial community^48^. In order to study this metabolic cross-feeding, they developed a pipeline where two GEMs are constrained to a minimal media condition, then iteratively simulated, thus updating the media with the costless metabolites until convergence. By FBA assumes that cells grow maximally and that all metabolites secretion enables maximum growth. Costless metabolites predicted to be secreted may differ for distinct feasible growth rates. Therefore, while the FBA-based perspective is valuable, it may only partially describe the simulated system. By allowing submaximal growth rates and alternative maxima through sampling, we demonstrate increased metabolic latitude for microbial communities.

A primary output from the costless secretion analysis is the number of iterations of model simulation until media convergence. We simulated 648 cases (pairwise combinations of 3 microbes, with and without oxygen, with 108 distinct fuel sources provided to supplement the minimal media). We assessed the number of iterations required to reach a steady media.

Interestingly, for both the normoxic and anoxic conditions, we see an increase in the number of rounds of model simulation with sampling compared to the base analysis with FBA. This makes sense, as a loosened restriction of growth rate allows for heterogeneous simulation results, which include a greater possible set of metabolites to be secreted and successive changes in the simulated media. That trend of more iterations with sampling remains even when we implement different cutoffs for whether a secreted metabolites is present in all, most, or at least one of the sets of sampled metabolic flux distributions (Figure 4A). Interestingly, the cutoff selected has much less of an effect than whether FBA (leftmost column) or flux sampling (right three columns) is chosen. We predict a much higher number of iterations in the anoxic condition, with up to 11 iterations of media change, compared to at most 3 rounds in the aerobic state.

**Figure 4:**
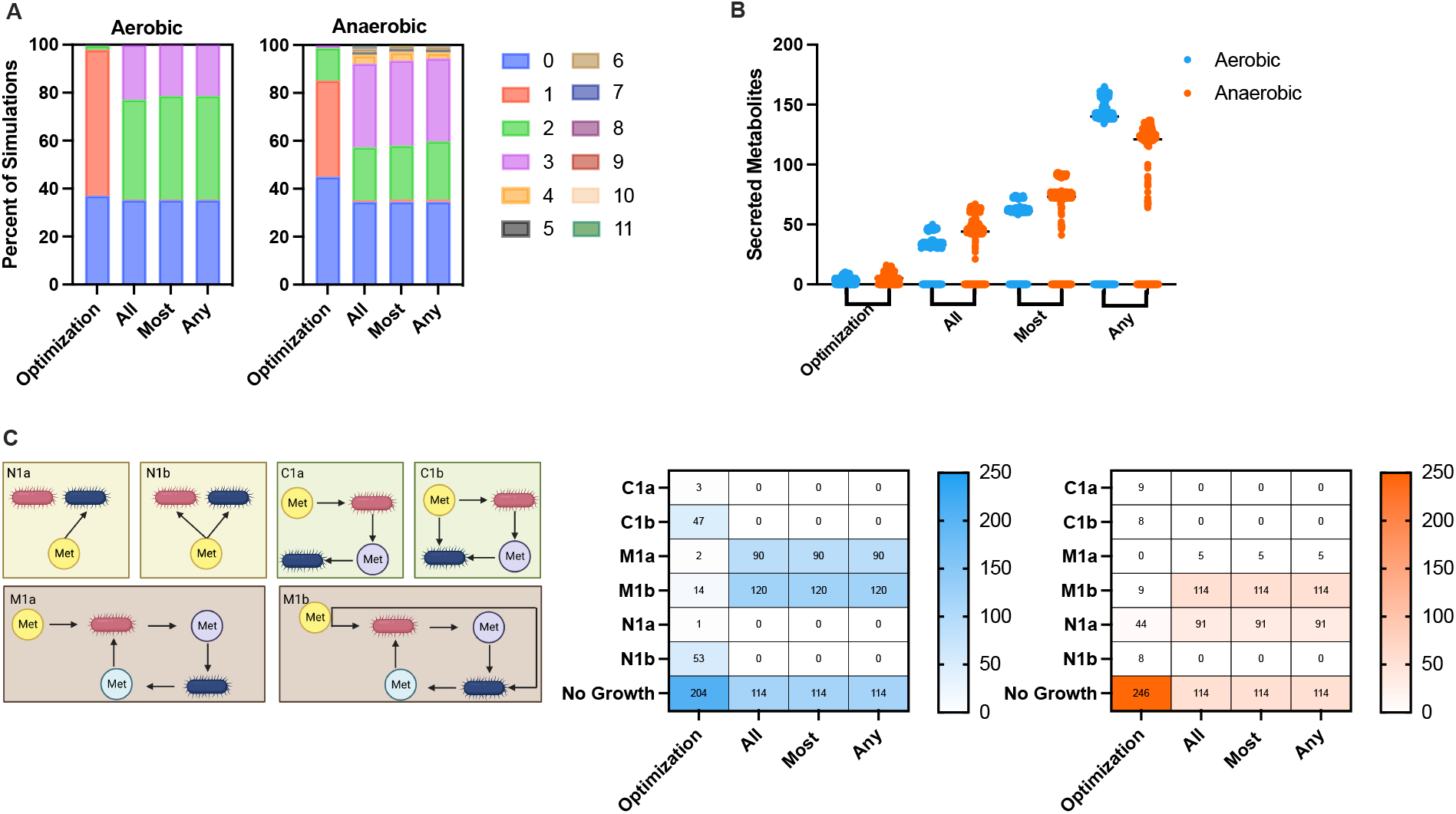
Costless secretion analysis. (A) Number of iterations required to achieve a stable media, for the aerobic (left) and anaerobic (right) states for optimization of biomass or different cutoffs for flux sampling (all, most, any), plotted as a percentage of all simulations. (B) Number of metabolites secreted for optimization of biomass or with distinct cutoffs for flux sampling (all, most, any) for anaerobic and aerobic conditions. (C) Each simulation was categorized into one of 7 cases (the six shown in the left panel and the case where no growth was achieved) for the aerobic or anaerobic condition.

We indeed see an increase in the number of metabolites secreted by the microbe pairs with flux sampling compared to FBA, as shown in Figure 4B. There is an apparent increase in predicted costless metabolites when the threshold is progressively loosened (from *all* to *most* to *any*). That is reasonable, as there are outliers or secreted metabolites that are particular to one or only few sampled flux distributions. We again predict an increase in secreted metabolites when simulating oxygen-free environments. Specifically, for anaerobic conditions, more unique metabolites are secreted as part of the cells’ metabolic flux patterns than in the oxygen-rich environments for the two most stringent cutoffs (*all* and *most*).

Pacheco established distinct interaction types, categorized based on the secretion and uptake rates of metabolites, using the following naming convention: non-interaction, or *N*, where no used media metabolites come from either model, commensalism, or *C*, characterized by unidirectional exchange, and mutualism, or *M* (where metabolites are interchanged between the two models). Following this letter designation, a numerical value represents the number of carbon sources added to the environment. Finally, the letters *a* or *b* specify the absence and presence of competition, respectively. As an example, N1a would describe a simulation where a metabolite is taken up by only one cell in the presence of one carbon source. We used the same naming convention to classify our simulations (Figure 4C). Firstly, we notice substantially fewer simulations with flux sampling where neither microbe achieved growth (204 compared to 114 of the 324 aerobic simulations and 246 compared to 114 of the 324 anaerobic simulations).

Though initially counterintuitive (that we would be more likely to achieve growth without optimizing for it than when attempting to maximize biomass), the result highlights the benefit of flux sampling. That is, due to the metabolic flexibility simulated with sampling, it is more likely that a microbe would secrete a metabolite beneficial for the other and thus enable the other cell to grow. In contrast, emergent interactive behavior is less likely when each microbe is “selfishly” oriented towards its growth at the expense of all other cellular goals (via FBA). We also note differences between the aerobic and anaerobic conditions, with the aerobic sampling simulations producing more instances of cooperative mutualism (M1a: 90 with aerobic sampling compared to 5 with anaerobic sampling) and the anaerobic simulations resulting in more non-competitive non-interaction (N1a: 91 with anaerobic sampling compared to 0 with aerobic sampling).

## 5. Discussion

Phenotypic heterogeneity, even in the monoculture of a genotypically uniform population, is known to have a substantial effect on observed community outcomes. However, the effects of this heterogeneity have yet to be fully studied, despite the rapid and substantial increases in modeling efforts at the genome-scale. In addition, microbes have been shown to exhibit sub-maximal growth, which needs to be sufficiently addressed with GEMs. While phenotypic heterogeneity and sub-maximal growth dynamics have been studied in individual GEMs of microbial activity, these two phenomena have not been analyzed for models of microbial interactions^30,52–58^. In this work, we demonstrate how pairing disparate existing approaches of flux sampling and modeling of communities pushes the field of metabolic modeling forward. We systematically evaluate the predictive effects of replacing FBA and its central assumption of maximal growth with flux sampling approaches.

In particular, we assess the effect of exploring the entire flux solution space with three distinct approaches of microbial community modeling: the compartmentalized approach, the lumped model or “enzyme soup” approach, and the costless secretion approach. With each approach, we replicate the major conclusions achieved with optimization of biomass using FBA For example, we predict higher frequency of cooperation under anaerobic conditions. Furthermore, applying flux sampling expands our understanding of the systems-level heterogeneity that gives rise to observed community activity. For the compartmentalized approach, we show increased tendency toward stable consortia and provide an ability to identify distinct growth rate-dependent interaction regimes. For the lumped modeling approach, we predict large differences in the predicted flux for certain pathways and reactions than others, and in the turnover of specific metabolites. With the costless secretion approach, we predict a substantially wider range of metabolites secreted, enabling growth on substrates that had not been predicted when optimizing biomass using FBA.

As previously found, most observable metabolic heterogeneity across a population has two primary sources: variation in network structure and variation in network usage (divergence in form and functional utilization)^59^. Ensemble modeling of GEMs has been shown to lead to increased accuracy and is of particular focus to the field with the emergence of novel tools; however, an equivalent effort has not been put towards understanding heterogeneous states achieved with a consistent network, despite the existence of flux sampling of GEMs as a tool for the past 20 years^60,61^. To our knowledge, one paper has used sampling to study cell-cell metabolic interactions^62^. Other researchers have identified this gap, and future work can more earnestly utilize and leverage the technique^63^.

We recognize some limitations of our work. A particular area for improvement of genome-scale modeling is the difficulty in assigning constraints for the reaction fluxes. Without appropriate bounds on metabolic reaction rates, flux sampling may explore biologically unreasonable metabolic states. The emergence of novel experimental tools is particularly promising to address this limitation. For example, -omics technologies enable *in vitro* and *in vivo* measurements of growth rates, metabolite secretion, and impact of enzymatic knockouts. Such data can be used to provide biologically reasonable constraints on reaction fluxes. In addition, we used thresholding to keep the analyses computationally feasible. However, this potentially limits our results. Improvements in computational ability, from advances in computing speed and algorithm development, will enable us to investigate the full range of biological outcomes possible with flux sampling without imposing artificial thresholds. Finally, we evaluated microbial fitness and interspecies relationships based on growth rate using flux sampling, eliminating the necessity of maximizing biomass. Future work can explore alternative metrics to assess cellular behavior. This is especially important because genome-scale modeling is increasingly used for eukaryotic (principally human) cells, where growth rate as a proxy for cell health is less supported^64–69^. For example, rather than focusing on growth, we could instead study flux through a specific reaction or pathway known to mediate the behavior of a particular cell type.

## 6. Conclusion

In this work, we evaluate the effect of flux sampling on three standard approaches for modeling the interactions between microbes at the genome-scale. The method clearly distinguishes between optimization-based and sampling-based characterizations of the metabolic interactions within a community. We demonstrate the utility of flux sampling in quantitatively studying metabolic interactions in microbial communities.

## 7. Acknowledgements

This work was supported by the National Cancer Institute of the National Institutes of Health grant 1U01CA232137 and administrative supplement (to S.D.F.). The authors thank members of the Finley research group for critical feedback.

## 8. Author contributions

P.E.G. and S.D.F. conceived of the presented idea. P.E.G. planned and carried out computational model development and simulations. S.D.F. supervised the project and provided financial support. All authors discussed the results and contributed to the final manuscript.

## Supplemental Information

**Figure S1:**
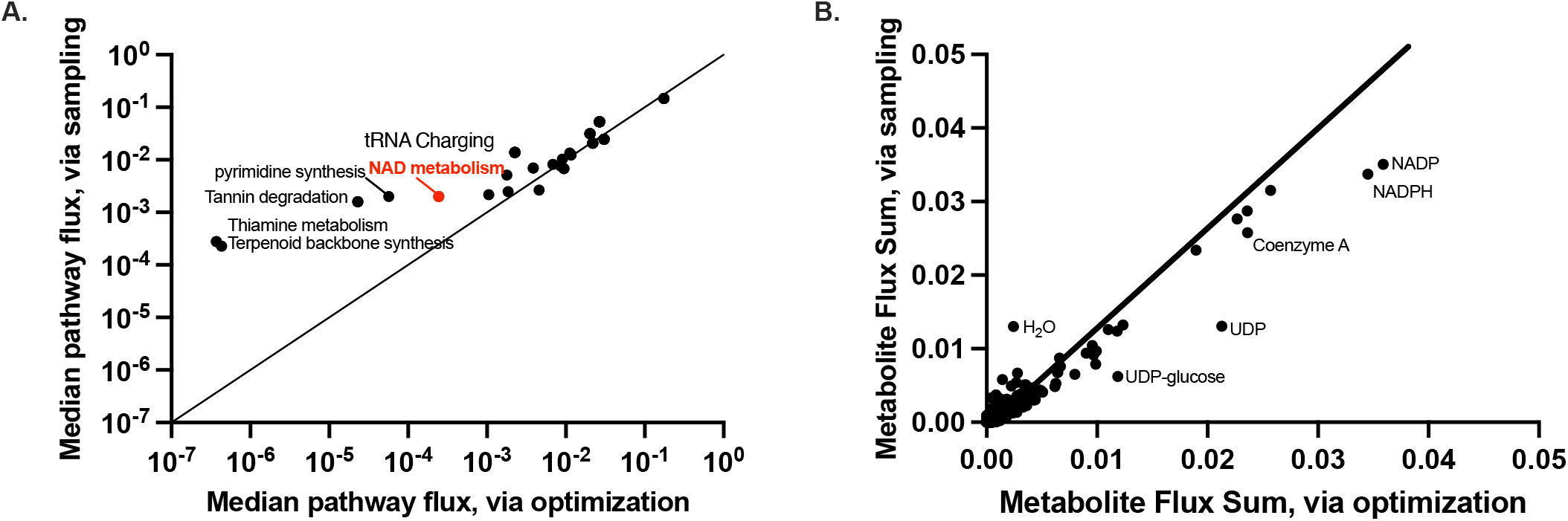
Community-constrained sampling compared to FBA optimization. **(A)** Median pathway flux values predicted by community-constrained flux sampling compared to optimization of biomass. Subsystems that have significantly different median fluxes are labeled. **(B)** Comparison of the flux-sum value for each metabolite for community-constrained flux sampling and optimization of biomass.

## References

1. Ponomarova, O., and Patil, K.R. (2015). Metabolic interactions in microbial communities: untangling the Gordian knot. Current Opinion in Microbiology 27, 37–44. 10.1016/j.mib.2015.06.014.

2. Douglas, A.E. (2020). The microbial exometabolome: ecological resource and architect of microbial communities. Philos Trans R Soc Lond B Biol Sci 375, 20190250. 10.1098/rstb.2019.0250.

3. Melkonian, C., Seidl, M.F., Hooft, J.J.J. van der, and Vos, M.G.J. de (2022). Metabolic interactions shape a community’s phenotype. Trends in Microbiology 30, 609–611. 10.1016/j.tim.2022.05.001.

4. Zhou, Z., Tran, P.Q., Breister, A.M., Liu, Y., Kieft, K., Cowley, E.S., Karaoz, U., and Anantharaman, K. (2022). METABOLIC: high-throughput profiling of microbial genomes for functional traits, metabolism, biogeochemistry, and community-scale functional networks. Microbiome 10, 33. 10.1186/s40168-021-01213-8.

5. Embree, M., Liu, J.K., Al-Bassam, M.M., and Zengler, K. (2015). Networks of energetic and metabolic interactions define dynamics in microbial communities. Proceedings of the National Academy of Sciences 112, 15450–15455. 10.1073/pnas.1506034112.

6. Khandelwal, R.A., Olivier, B.G., Röling, W.F.M., Teusink, B., and Bruggeman, F.J. (2013). Community Flux Balance Analysis for Microbial Consortia at Balanced Growth. PLoS One 8, e64567. 10.1371/journal.pone.0064567.

7. Tzamali, E., Poirazi, P., Tollis, I.G., and Reczko, M. (2011). A computational exploration of bacterial metabolic diversity identifying metabolic interactions and growth-efficient strain communities. BMC Syst Biol 5, 167. 10.1186/1752-0509-5-167.

8. Gu, C., Kim, G.B., Kim, W.J., Kim, H.U., and Lee, S.Y. (2019). Current status and applications of genome-scale metabolic models. Genome Biology 20, 121. 10.1186/s13059-019-1730-3.

9. Sertbas, M., and Ulgen, K.O. (2020). Genome-Scale Metabolic Modeling for Unraveling Molecular Mechanisms of High Threat Pathogens. Frontiers in Cell and Developmental Biology 8.

10. Zhang, C., Qi, J., and Cao, Y. (2014). Synergistic Effect of Yeast-Bacterial Co-Culture on Bioremediation of Oil-Contaminated Soil. Bioremediation Journal 18, 136–146. 10.1080/10889868.2013.847402.

11. Thommes, M., Wang, T., Zhao, Q., Paschalidis, I.C., and Segrè, D. (2019). Designing Metabolic Division of Labor in Microbial Communities. mSystems 4, e00263–18. 10.1128/mSystems.00263-18.

12. Reed, J.L. (2017). Genome-Scale Metabolic Modeling and Its Application to Microbial Communities (National Academies Press (US)).

13. Cuevas, D.A., Edirisinghe, J., Henry, C.S., Overbeek, R., O’Connell, T.G., and Edwards, R.A. (2016). From DNA to FBA: How to Build Your Own Genome-Scale Metabolic Model. Frontiers in Microbiology 7.

14. Proffitt, C., Bidkhori, G., Lee, S., Tebani, A., Mardinoglu, A., Uhlen, M., Moyes, D.L., and Shoaie, S. (2022). Genome-scale metabolic modelling of the human gut microbiome reveals changes in the glyoxylate and dicarboxylate metabolism in metabolic disorders. iScience 25, 104513. 10.1016/j.isci.2022.104513.

15. Ankrah, N.Y.D., Bernstein, D.B., Biggs, M., Carey, M., Engevik, M., García-Jiménez, B., Lakshmanan, M., Pacheco, A.R., Sulheim, S., and Medlock, G.L. (2021). Enhancing Microbiome Research through Genome-Scale Metabolic Modeling. mSystems 6, e00599–21. 10.1128/mSystems.00599-21.

16. Zorrilla, F., Buric, F., Patil, K.R., and Zelezniak, A. (2021). metaGEM: reconstruction of genome scale metabolic models directly from metagenomes. Nucleic Acids Research 49, e126. 10.1093/nar/gkab815.

17. Magnúsdóttir, S., Heinken, A., Kutt, L., Ravcheev, D.A., Bauer, E., Noronha, A., Greenhalgh, K., Jäger, C., Baginska, J., Wilmes, P., et al. (2017). Generation of genome-scale metabolic reconstructions for 773 members of the human gut microbiota. Nat Biotechnol 35, 81–89. 10.1038/nbt.3703.

18. Ang, K.S., Lakshmanan, M., Lee, N.-R., and Lee, D.-Y. (2018). Metabolic Modeling of Microbial Community Interactions for Health, Environmental and Biotechnological Applications. Curr Genomics 19, 712–722. 10.2174/1389202919666180911144055.

19. Sonawane, J.M., Mahadevan, R., Pandey, A., and Greener, J. (2022). Recent progress in microbial fuel cells using substrates from diverse sources. Heliyon 8, e12353. 10.1016/j.heliyon.2022.e12353.

20. Frioux, C., Singh, D., Korcsmaros, T., and Hildebrand, F. (2020). From bag-of-genes to bag-of-genomes: metabolic modelling of communities in the era of metagenome-assembled genomes. Comput Struct Biotechnol J 18, 1722–1734. 10.1016/j.csbj.2020.06.028.

21. R. Dillard, L. D. Payne D., and A. Papin J. (2021). Mechanistic models of microbial community metabolism. Molecular Omics 17, 365–375. 10.1039/D0MO00154F.

22. Roume, H., Heintz-Buschart, A., Muller, E.E.L., May, P., Satagopam, V.P., Laczny, C.C., Narayanasamy, S., Lebrun, L.A., Hoopmann, M.R., Schupp, J.M., et al. (2015). Comparative integrated omics: identification of key functionalities in microbial community-wide metabolic networks. npj Biofilms Microbiomes 1, 1–11. 10.1038/npjbiofilms.2015.7.

23. Faria, J.P., Khazaei, T., Edirisinghe, J.N., Weisenhorn, P., Seaver, S.D., Conrad, N., Harris, N., DeJongh, M., and Henry, C.S. (2016). Constructing and Analyzing Metabolic Flux Models of Microbial Communities. Springer Protocols Handbooks Hydrocarbon and Lipid Microbiology Protocols. 10.1007/8623_2016_215.

24. Heinken, A., and Thiele, I. (2015). Anoxic Conditions Promote Species-Specific Mutualism between Gut Microbes In Silico. Applied and Environmental Microbiology 81, 4049–4061. 10.1128/AEM.00101-15.

25. Stolyar, S., Van Dien, S., Hillesland, K.L., Pinel, N., Lie, T.J., Leigh, J.A., and Stahl, D.A. (2007). Metabolic modeling of a mutualistic microbial community. Molecular Systems Biology 3, 92. 10.1038/msb4100131.

26. Wintermute, E.H., and Silver, P.A. (2010). Emergent cooperation in microbial metabolism. Molecular Systems Biology 6, 407. 10.1038/msb.2010.66.

27. Blasco, T., Pérez-Burillo, S., Balzerani, F., Hinojosa-Nogueira, D., Lerma-Aguilera, A., Pastoriza, S., Cendoya, X., Rubio, Á., Gosalbes, M.J., Jiménez-Hernández, N., et al. (2021). An extended reconstruction of human gut microbiota metabolism of dietary compounds. Nat Commun 12, 4728. 10.1038/s41467-021-25056-x.

28. Orth, J.D., Thiele, I., and Palsson, B.Ø.xs (2010). What is flux balance analysis? Nature Biotechnology 28, 245–248. 10.1038/nbt.1614.

29. Celiker, H., and Gore, J. (2012). Competition between species can stabilize public-goods cooperation within a species. Mol Syst Biol 8, 621. 10.1038/msb.2012.54.

30. Raman, K., and Chandra, N. (2009). Flux balance analysis of biological systems: applications and challenges. Briefings in Bioinformatics 10, 435–449. 10.1093/bib/bbp011.

31. Diener, C., and Gibbons, S.M. (2023). More is Different: Metabolic Modeling of Diverse Microbial Communities. mSystems, e01270–22. 10.1128/msystems.01270-22.

32. García Sánchez, C.E., and Torres Sáez, R.G. (2014). Comparison and analysis of objective functions in flux balance analysis. Biotechnology Progress 30, 985–991. 10.1002/btpr.1949.

33. Schnitzer, B., Österberg, L., and Cvijovic, M. (2022). The choice of the objective function in flux balance analysis is crucial for predicting replicative lifespans in yeast. PLOS ONE 17, e0276112. 10.1371/journal.pone.0276112.

34. Lachance, J.-C., Lloyd, C.J., Monk, J.M., Yang, L., Sastry, A.V., Seif, Y., Palsson, B.O., Rodrigue, S., Feist, A.M., King, Z.A., et al. (2019). BOFdat: Generating biomass objective functions for genome-scale metabolic models from experimental data. PLOS Computational Biology 15, e1006971. 10.1371/journal.pcbi.1006971.

35. Feist, A.M., and Palsson, B.O. (2010). The Biomass Objective Function. Curr Opin Microbiol 13, 344–349. 10.1016/j.mib.2010.03.003.

36. Herrmann, H.A., Dyson, B.C., Vass, L., Johnson, G.N., and Schwartz, J.-M. (2019). Flux sampling is a powerful tool to study metabolism under changing environmental conditions. npj Syst Biol Appl 5, 1–8. 10.1038/s41540-019-0109-0.

37. Bordel, S., Agren, R., and Nielsen, J. (2010). Sampling the Solution Space in Genome-Scale Metabolic Networks Reveals Transcriptional Regulation in Key Enzymes. PLOS Computational Biology 6, e1000859. 10.1371/journal.pcbi.1000859.

38. Fallahi, S., Skaug, H.J., and Alendal, G. (2020). A comparison of Monte Carlo sampling methods for metabolic network models. PLOS ONE 15, e0235393. 10.1371/journal.pone.0235393.

39. Scott, W.T., Smid, E.J., Block, D.E., and Notebaart, R.A. (2021). Metabolic flux sampling predicts strain-dependent differences related to aroma production among commercial wine yeasts. Microbial Cell Factories 20, 204. 10.1186/s12934-021-01694-0.

40. Martino, D.D., Mori, M., and Parisi, V. (2015). Uniform Sampling of Steady States in Metabolic Networks: Heterogeneous Scales and Rounding. PLOS ONE 10, e0122670. 10.1371/journal.pone.0122670.

41. Heinken, A., Hertel, J., Acharya, G., Ravcheev, D.A., Nyga, M., Okpala, O.E., Hogan, M., Magnúsdóttir, S., Martinelli, F., Nap, B., et al. (2023). Genome-scale metabolic reconstruction of 7,302 human microorganisms for personalized medicine. Nat Biotechnol, 1–12. 10.1038/s41587-022-01628-0.

42. Klitgord, N., and Segrè, D. (2010). Environments that induce synthetic microbial ecosystems. PLoS Comput Biol 6, e1001002. 10.1371/journal.pcbi.1001002.

43. Kook, Y., Lee, Y.T., Shen, R., and Vempala, S.S. (2022). Sampling with Riemannian Hamiltonian Monte Carlo in a Constrained Space.

44. Eng, A., and Borenstein, E. (2016). An algorithm for designing minimal microbial communities with desired metabolic capacities. Bioinformatics 32, 2008–2016. 10.1093/bioinformatics/btw107.

45. Frioux, C., Fremy, E., Trottier, C., and Siegel, A. (2018). Scalable and exhaustive screening of metabolic functions carried out by microbial consortia. Bioinformatics 34, i934–i943. 10.1093/bioinformatics/bty588.

46. Greenblum, S., Turnbaugh, P.J., and Borenstein, E. (2012). Metagenomic systems biology of the human gut microbiome reveals topological shifts associated with obesity and inflammatory bowel disease. Proc Natl Acad Sci U S A 109, 594–599. 10.1073/pnas.1116053109.

47. Ofaim, S., Ofek-Lalzar, M., Sela, N., Jinag, J., Kashi, Y., Minz, D., and Freilich, S. (2017). Analysis of Microbial Functions in the Rhizosphere Using a Metabolic-Network Based Framework for Metagenomics Interpretation. Front Microbiol 8, 1606. 10.3389/fmicb.2017.01606.

48. Pacheco, A.R., Moel, M., and Segrè, D. (2019). Costless metabolic secretions as drivers of interspecies interactions in microbial ecosystems. Nat Commun 10, 103. 10.1038/s41467-018-07946-9.

49. Blanchard, A.E., and Lu, T. (2015). Bacterial social interactions drive the emergence of differential spatial colony structures. BMC Systems Biology 9, 59. 10.1186/s12918-015-0188-5.

50. Boza, G., Barabás, G., Scheuring, I., and Zachar, I. (2023). Eco-evolutionary modelling of microbial syntrophy indicates the robustness of cross-feeding over cross-facilitation. Sci Rep 13, 907. 10.1038/s41598-023-27421-w.

51. Kullback, S., and Leibler, R.A. (1951). On Information and Sufficiency. The Annals of Mathematical Statistics 22, 79–86. 10.1214/aoms/1177729694.

52. Chung, C.H., and Chandrasekaran, S. (2022). A flux-based machine learning model to simulate the impact of pathogen metabolic heterogeneity on drug interactions. PNAS Nexus 1, pgac132. 10.1093/pnasnexus/pgac132.

53. Damiani, C., Maspero, D., Filippo, M.D., Colombo, R., Pescini, D., Graudenzi, A., Westerhoff, H.V., Alberghina, L., Vanoni, M., and Mauri, G. (2019). Integration of single-cell RNA-seq data into population models to characterize cancer metabolism. PLOS Computational Biology 15, e1006733. 10.1371/journal.pcbi.1006733.

54. Zampieri, G., Vijayakumar, S., Yaneske, E., and Angione, C. (2019). Machine and deep learning meet genome-scale metabolic modeling. PLOS Computational Biology 15, e1007084. 10.1371/journal.pcbi.1007084.

55. Øyås, O., and Stelling, J. (2018). Genome-scale metabolic networks in time and space. Current Opinion in Systems Biology 8, 51–58. 10.1016/j.coisb.2017.12.003.

56. Jouhten, P., Wiebe, M., and Penttilä, M. (2012). Dynamic flux balance analysis of the metabolism of Saccharomyces cerevisiae during the shift from fully respirative or respirofermentative metabolic states to anaerobiosis. The FEBS Journal 279, 3338–3354. 10.1111/j.1742-4658.2012.08649.x.

57. Diener, C., Gibbons, S.M., and Resendis-Antonio, O. (2020). MICOM: Metagenome-Scale Modeling To Infer Metabolic Interactions in the Gut Microbiota. mSystems 5, e00606–19. 10.1128/mSystems.00606-19.

58. Panikov, N.S. (2021). Genome-Scale Reconstruction of Microbial Dynamic Phenotype: Successes and Challenges. Microorganisms 9, 2352. 10.3390/microorganisms9112352.

59. Cabbia, A., Hilbers, P.A.J., and van Riel, N.A.W. (2020). A Distance-Based Framework for the Characterization of Metabolic Heterogeneity in Large Sets of Genome-Scale Metabolic Models. Patterns (N Y) 1, 100080. 10.1016/j.patter.2020.100080.

60. Medlock, G.L., Moutinho, T.J., and Papin, J.A. (2020). Medusa: Software to build and analyze ensembles of genome-scale metabolic network reconstructions. PLOS Computational Biology 16, e1007847. 10.1371/journal.pcbi.1007847.

61. Biggs, M.B., and Papin, J.A. (2017). Managing uncertainty in metabolic network structure and improving predictions using EnsembleFBA. PLOS Computational Biology 13, e1005413. 10.1371/journal.pcbi.1005413.

62. Benito-Vaquerizo, S., Diender, M., Parera Olm, I., Martins dos Santos, V.A.P., Schaap, P.J., Sousa, D.Z., and Suarez-Diez, M. (2020). Modeling a co-culture of Clostridium autoethanogenum and Clostridium kluyveri to increase syngas conversion to medium-chain fatty-acids. Comput Struct Biotechnol J 18, 3255–3266. 10.1016/j.csbj.2020.10.003.

63. Scott, W.T., Benito-Vaquerizo, S., Zimmerman, J., Bajić, D., Heinken, A., Suarez-Diez, M., and Schaap, P.J. (2023). A structured evaluation of genome-scale constraint-based modeling tools for microbial consortia. 2023.02.08.527721. 10.1101/2023.02.08.527721.

64. Wang, H., Robinson, J.L., Kocabas, P., Gustafsson, J., Anton, M., Cholley, P.-E., Huang, S., Gobom, J., Svensson, T., Uhlen, M., et al. (2021). Genome-scale metabolic network reconstruction of model animals as a platform for translational research. Proceedings of the National Academy of Sciences 118, e2102344118. 10.1073/pnas.2102344118.

65. Nilsson, A., and Nielsen, J. (2017). Genome scale metabolic modeling of cancer. Metab Eng 43, 103–112. 10.1016/j.ymben.2016.10.022.

66. Wang, J., Delfarah, A., Gelbach, P.E., Fong, E., Macklin, P., Mumenthaler, S.M., Graham, N.A., and Finley, S.D. (2022). Elucidating tumor-stromal metabolic crosstalk in colorectal cancer through integration of constraint-based models and LC-MS metabolomics. Metab Eng 69, 175–187. 10.1016/j.ymben.2021.11.006.

67. Gelbach, P.E., and Finley, S.D. (2023). Ensemble-based genome-scale modeling predicts metabolic differences between macrophage subtypes in colorectal cancer. 2023.03.09.532000. 10.1101/2023.03.09.532000.

68. Frades, I., Foguet, C., Cascante, M., and Araúzo-Bravo, M.J. (2021). Genome Scale Modeling to Study the Metabolic Competition between Cells in the Tumor Microenvironment. Cancers (Basel) 13, 4609. 10.3390/cancers13184609.

69. Brunk, E., Sahoo, S., Zielinski, D.C., Altunkaya, A., Dräger, A., Mih, N., Gatto, F., Nilsson, A., Preciat Gonzalez, G.A., Aurich, M.K., et al. (2018). Recon3D enables a three-dimensional view of gene variation in human metabolism. Nat Biotechnol 36, 272–281. 10.1038/nbt.4072.

